# ComplexDesign: sequence-hallucination design of protein binders bridging multiple proteins

**DOI:** 10.64898/2026.06.21.733655

**Authors:** Jing Xu, Milong Ren, Ning Qi, Xinru Zhang, Zaikai He, Chungong Yu, Dongbo Bu

**Affiliations:** SKLP, Institute of Computing Technology, Chinese Academy of Sciences, Beijing, 100190, China; School of Advanced Interdisciplinary Sciences, University of Chinese Academy of Sciences, Beijing, 100049, China; University of Chinese Academy of Sciences, Beijing, 100049, China; Central China Institute of Artificial Intelligence, Zhengzhou, Henan, 450046, China

## Abstract

**Motivation:** Designing multichain protein complexes requires coordinating the folding of component proteins with the formation of their interfaces. The existing methods, however, remain limited in their ability to satisfy these requirements simultaneously, especially for trimeric and tetrameric complexes. As an important practical scenario, designing a binder that bridges two target proteins into a ternary complex requires flexibility in the relative arrangement of the two targets, adding an additional challenge to existing design methods.

**Results:** We present ComplexDesign, a hallucination-based approach for multichain protein design. ComplexDesign performs structure-prediction-guided sequence optimization to simultaneously fold each protein chain and form inter-chain interactions that bind them together. To provide the flexibility required to appropriately arrange these target proteins, ComplexDesign introduces a specialized masking mechanism that enables exploration of possible relative arrangements rather than being limited to the predefined ones. Across a comprehensive set of benchmarks with various chain lengths, ComplexDesign outperformed existing methods in the unconditional design of dimers, trimers, and tetramers, achieving a high design success rate exceeding 50%, supporting its capability for multichain complex design. Furthermore, in the case of multi-target binder design, ComplexDesign produced high-confidence, self-consistent ternary complexes for 8 out of 10 target pairs. These results establish ComplexDesign as an effective tool for multichain protein design, with particular utility for designing binders that bridge two target proteins.

**Availability and implementation:** The source code of ComplexDesign will be made publicly available upon publication.

## 1. Introduction

Protein complexes are central to many biological processes and applications, making their design an important goal in computational biology (Greenblatt et al., 2024). Unlike the design of single-chain proteins, protein complex design should preserve the structural integrity of each component protein while also forming the inter-chain interactions to bind them together (Zhu et al., 2021). Recent advances in machine learning have enabled the design of increasingly complex protein structures and assemblies (Listov et al., 2024; Albanese et al., 2025).

The existing methods for protein complex design fall broadly into two categories: hallucination-based optimization and training-based generative modeling. In hallucination-based approaches, sequences are iteratively optimized by a structure prediction model to find a sequence whose predicted structure possesses the intended properties (Anishchenko et al., 2021). For instance, RSO (Frank et al., 2024) applied this strategy in a relaxed sequence space for scalable protein design. BindCraft (Pacesa et al., 2025) extended the same general principle to automatic *de novo* binder design and showed experimental success across diverse targets. HalluDesign (Fang et al., 2025) further extended this line of work with a forward-pass-only framework for iterative sequence-structure co-optimization. In parallel, the training-based methods employ a different strategy: they learn distributions from structural data and sample proteins or complexes directly from these distributions. These methods can operate at the backbone level, say diffusion-based RFdiffusion (Watson et al., 2023) and the programmable generative model Chroma (Ingraham et al., 2023), or at the all-atom level, say APM for multichain complex generation (Chen et al., 2025), as well as Latent-X (Latent Labs Team, 2025), RFdiffusion3 (Butcher et al., 2025), and BoltzGen (Stark et al., 2025) for binder and biomolecular interaction design.

Despite these promising progresses, the existing approaches still have limitations in broader multichain protein design settings: In the case of unconditional multichain design, existing generative methods, such as APM and Chroma, do not yet reliably produce complexes that satisfy both chain-level structural integrity and complex-level inter-chain organization, especially for higher-order assemblies such as trimers and tetramers.

A more challenging task arises from binder design: current methods are typically designed for a single target, such as a structure, binding site, or epitope (Butcher et al., 2025; Pacesa et al., 2025; Latent Labs Team, 2025; Stark et al., 2025). Even when the input target contains multiple chains, these methods generally treat them as a single preassembled target for binder design, rather than exploring a binder against two distinct targets with uncertain relative positions. As a result, these methods do not directly address multi-target binder design, in which a single binder must engage multiple distinct targets. In this study, we focus on a practically important scenario with two targets: designing a single *de novo* binder that bridges two target proteins to form a ternary complex. These ternary assemblies have significant application in a range of cases, including molecular glues, PROTACs, bioPROTACs, AbTACs, and bispecific antibody-based therapies (Sasso et al., 2022; Békés et al., 2022; Shen and Dassama, 2023; Cotton et al., 2021; Klein et al., 2024).

To address these challenges, we propose ComplexDesign, a sequence-hallucination framework for designing multichain protein complexes, with a particular focus on binders that bridge two target proteins. ComplexDesign performs structure-prediction-guided sequence optimization by backpropagating objective gradients to designable sequence representations. This approach enables simultaneous optimization of intra-chain folding and inter-chain interface formation for multichain complexes. To design multi-target binders, ComplexDesign introduces a masking mechanism that removes target–target pairwise information during optimization. This mechanism prevents the two targets from being imposed as a fixed joint template, thus providing flexibility in their relative arrangement. The binder is therefore optimized to bind both target proteins, allowing the two binder– target interfaces and the final ternary complex to emerge during design.

We first evaluate ComplexDesign on unconditional multichain design benchmarks. Across dimer, trimer, and tetramer settings, ComplexDesign outperformed representative generative baseline methods and achieved success rates exceeding 50% for all evaluated chain lengths. These results demonstrate ComplexDesign’s capacity to design multichain complexes. We then evaluate ComplexDesign on a multi-target binder-design benchmark derived from MG-PDB (Liao et al., 2025). In this case, ComplexDesign yielded self-consistent ternary designs with scRMSD ≤ 2.0 Å for 8 out of 10 target pairs, with considerably high AF3 ipTM exceeding 0.85. These results establish ComplexDesign as an effective computational tool for multichain protein design, particularly for designing binders that bridge two separate targets to form ternary complexes.

## 2. Methods

### 2.1. Overview of ComplexDesign

ComplexDesign is a sequence-hallucination-based framework for multichain protein complex design (Fig. 1). The workflow consists of three main components, including structure-prediction-guided sequence optimization, inverse-folding-based sequence redesign, and structure-prediction-based evaluation, which are described below.

**Fig. 1:**
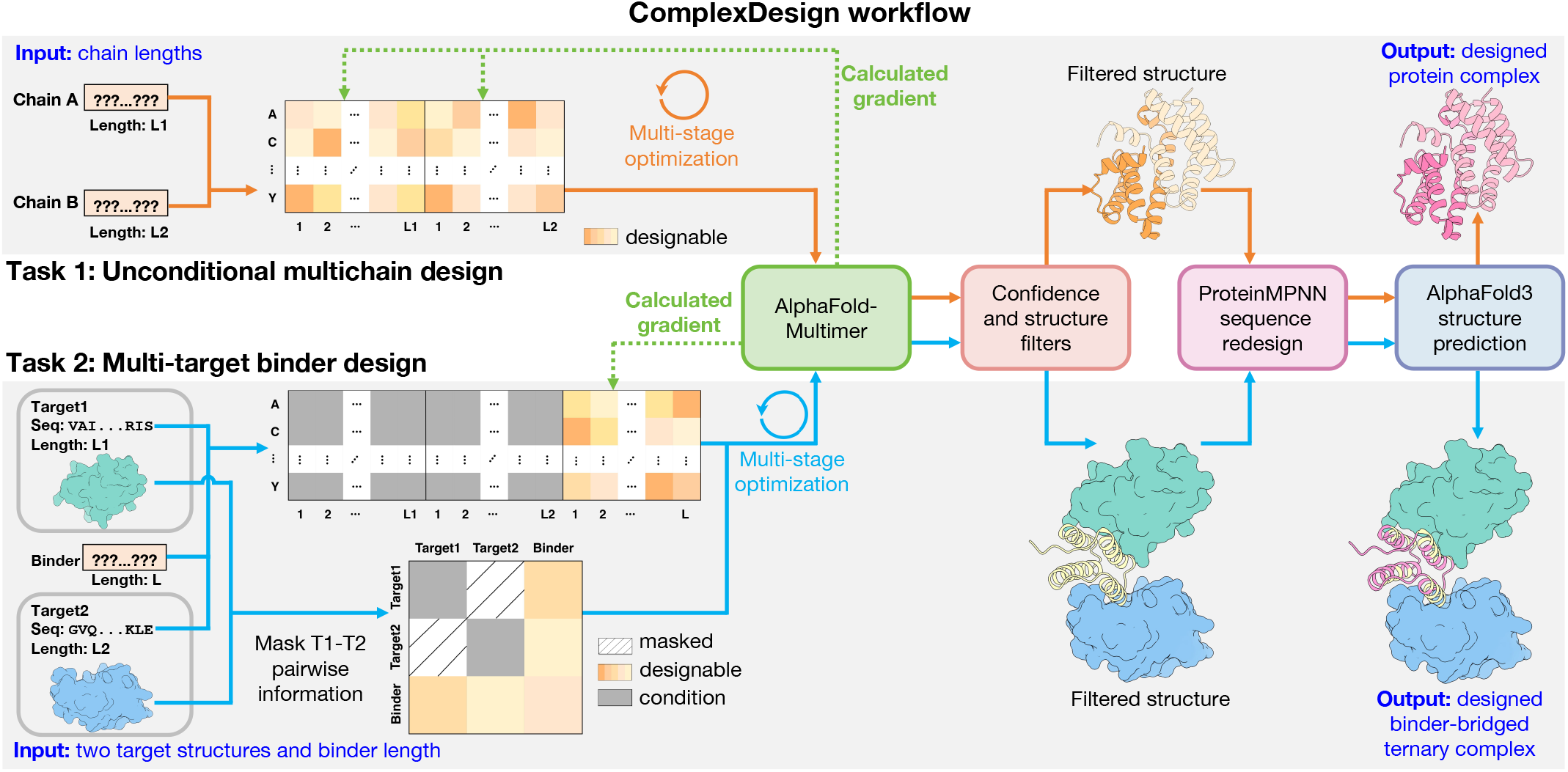
Overview of ComplexDesign. ComplexDesign uses a unified optimization–redesign–evaluation workflow for unconditional multichain design and multi-target binder design. In unconditional design, the input specifies only chain lengths. All chains are optimized jointly. In a multi-target binder design, a binder length is specified, and two target structures are provided as separate structural templates. The target sequences are kept fixed, whereas only the binder sequence is optimized. Target–target pairwise information is masked to avoid imposing a fixed target–target arrangement. In both tasks, AlphaFold-Multimer guides multi-stage sequence optimization by backpropagating objective gradients to the designable sequence representations. Designs that pass confidence and structural filters are then redesigned with ProteinMPNN and evaluated with AlphaFold3. **Alt text:** Schematic of the ComplexDesign workflow for two protein-complex design tasks. In unconditional multichain design, the input specifies the chain lengths, and sequence representations for all chains are optimized jointly. In multi-target binder design, the input comprises two target structures and a binder length; the target sequences remain fixed, whereas only the binder sequence representation is optimized. Target-target pairwise information is masked to avoid imposing a fixed relative arrangement between the two targets. In both tasks, AlphaFold-Multimer predicts the complex and backpropagates objective gradients to the designable sequence representations, followed by confidence and structural filtering, ProteinMPNN sequence redesign, and AlphaFold3 evaluation.

1. The sequence optimization module uses AlphaFold-Multimer (Evans et al., 2021) as the structure predictor, and the designable sequence representations are iteratively updated by backpropagating objective gradients from AlphaFold-Multimer predictions.
2. After sequence optimization, the designs that satisfy the confidence and structure filtering criteria are redesigned using ProteinMPNN (Dauparas et al., 2022) to improve sequence realizability.
3. Finally, the redesigned sequences are evaluated with AlphaFold3 (Abramson et al., 2024) to assess structural confidence, self-consistency, and interface quality.

ComplexDesign can accomplish two types of protein complex design tasks, i.e., unconditional multichain design and multi-target binder design, with the former providing the foundation for the latter. The two tasks use the same optimization framework but differ in their inputs, designable regions, and objective functions.

i. *Unconditional multichain design:* In this case, the input specifies only the number of chains and their lengths, and no structural template or fixed sequence is provided. All residues of the chains are designable, and the sequence optimization is performed jointly over all chains. Chain identities are explicitly encoded to enable simultaneous optimization of intra-chain folding and inter-chain interface formation.
ii. *Multi-target binder design:* Here, we focus on the application-relevant two-target setting, where a single *de novo* binder is designed to bridge two target proteins into a ternary complex. The input consists of a binder-length range and structures of the two targets. In each design run, a binder length is sampled from the specified range. The target sequences are fixed during optimization, and only the binder sequence is designable. To avoid constraining the design to a predefined target–target arrangement, ComplexDesign removes target–target pairwise information during optimization. Specifically, relative positional encoding and inter-chain template information between the two targets are masked. The binder is therefore optimized to bind to the two target proteins; thus, the binder–target interfaces and ternary geometry are determined during optimization rather than inherited from the input structures. Notably, by varying chain lengths and random seeds, ComplexDesign can yield complexes with different relative arrangements of the target chains, thereby providing diversity in the designed complexes.

### 2.2. Structure-prediction-guided sequence optimization

#### 2.2.1. Sequence representation and staged optimization

ComplexDesign optimizes protein complex sequences directly in sequence space using AlphaFold-Multimer (Evans et al., 2021) as the structure predictor to provide optimization signals. Here, the designable residues are represented as residue logits. ComplexDesign begins with these logits initialized with small random noise, and iterates multiple steps to optimize the sequence representations. At each iteration step, the current sequence representation is fed into AlphaFold-Multimer to predict a multichain structure. Next, the optimization objective value is computed from the predicted structure, and the corresponding gradients with respect to the sequence logits are backpropagated to update the sequence representation.

In particular, the sequence optimization consists of four stages, including *logits, softmax, one-hot*, and *semi-greedy*, as performed by ColabDesign and BindCraft (Ovchinnikov et al., 2025; Pacesa et al., 2025). This four-stage scheme progressively converts a continuous sequence representation into a discrete designed sequence. The *logits* and *softmax* stages optimize continuous and probability-based representations of sequences. The *one-hot* stage further sharpens the sequence representation toward discrete residue assignments. The final *semi-greedy* stage refines the current design through discrete mutation-based search.

#### 2.2.2. The objective function of sequence optimization

At each iteration, the predicted complex is evaluated by a weighted-sum objective function:

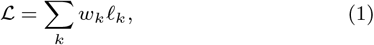

where *ℓ*_*k*_ denotes an individual loss term and *w*_*k*_ denotes its weight. The objective includes AF-derived confidence terms (pLDDT and i pTM), predicted-alignment-error terms (pAE and i pAE), and contact-based terms (contact and i contact). Terms evaluated within designable chains promote the structural integrity of the designed components, whereas interface-specific terms aim to promote compatible inter-chain contacts and relative placement. The details of these terms are provided in the Supplementary Material.

Notably, these terms are evaluated over different residue sets in the two ComplexDesign tasks:

i. In the case of unconditional multichain design, the pLDDT loss is applied to all residues. The pAE and contact loss are computed over the same-chain residue pairs, whereas the i pAE, i contact, and i pTM losses are computed using different-chain residue pairs. The objective includes pLDDT, pAE, contact, i pAE, i contact, and i pTM losses, with weights of 0.1, 0.4, 1.0, 0.1, 1.0, and 0.05, respectively.
ii. In the case of multi-target binder design, the same loss terms are used with the same weights, but evaluation is primarily restricted to the binder and the two binder–target interfaces. Specifically, the pLDDT and contact losses are computed within the binder. The pAE loss is computed for residue pairs that involve the binder. The i pAE and i contact losses are applied to binder–target residue pairs, whereas the i pTM loss is computed using all different-chain residue pairs in the ternary complex. In addition, a radius-of-gyration loss with weight 0.3 is applied to the binder. This term discourages overly extended conformations and promotes compact folded states.

### 2.3. Filtering and evaluating the designed complexes

ComplexDesign filters the complexes obtained from the four-stage optimization procedure based on their predicted structures. The filtered structures are then redesigned with ProteinMPNN before final structure-prediction-based evaluation. The details of these steps are described below.

Filtering is applied during and after optimization. At the end of each stage, the lowest-loss structure predicted in that stage is considered. Optimization proceeds to the next stage only if its mean pLDDT over the designable residues exceeds 0.65. After all four stages are completed, the final structure is retained only if it again meets this pLDDT threshold and contains no C*α* clashes. A C*α* clash is defined as a pair of C*α* atoms closer than 2.5 Å, excluding adjacent residues within the same chain. For multi-target binder design, the final predicted ternary structure must also contain at least three binder residues in contact with either target chain. A binder residue is defined as an interface residue if any of its atoms lie within 3.5 Å of either target chain.

Filtered structures are then passed to ProteinMPNN (Dauparas et al., 2022) for sequence redesign. In the unconditional design setting, all chains are redesigned. In the multi-target binder setting, only non-interface binder residues are redesigned, whereas binder residues at the designed interfaces are kept fixed to preserve the designed interactions.

Finally, redesigned sequences are re-predicted with AlphaFold3 (AF3), a high-accuracy model for complex-structure prediction (Abramson et al., 2024). The resulting predicted structures are then used to assess structural confidence, structural recovery, and interface quality after sequence redesign.

## 3. Results

We first evaluated ComplexDesign’s performance in unconditional multichain design, as in prior studies (Ingraham et al., 2023; Chen et al., 2025). Next, we evaluated ComplexDesign on a multi-target binder design benchmark derived from MG-PDB (Liao et al., 2025). This benchmark comprised ten target pairs spanning a range of difficulty levels, biological contexts, and practical relevance. Across both tasks, we used AlphaFold3 confidence scores, self-consistency, and Rosetta-based interface energy as metrics.

### 3.1. Data sets and evaluation criteria

#### 3.1.1. Unconditional multichain design

We evaluated ComplexDesign across nine chain-length combinations spanning dimer, trimer, and tetramer settings. For each setting, we designed 32 complexes and generated eight ProteinMPNN redesigns per complex. We compared ComplexDesign with two representative baseline methods, APM (Chen et al., 2025) and Chroma (Ingraham et al., 2023), under the same sampling settings. All generated complexes and their ProteinMPNN redesigns were folded with AlphaFold3 in single-sequence mode, without template or MSA information for any chain.

Across all settings, we used overall AlphaFold3 ipTM and C*α* self-consistency RMSD (scRMSD) (Bennett et al., 2023) as the main evaluation metrics. We summarized success rates using thresholds of overall ipTM ≥ 0.8 and scRMSD ≤ 2.0 Å. These cutoffs correspond to confident high-quality complex predictions (Abramson et al., 2024) and stringent structural self-consistency (Bennett et al., 2023; Frank et al., 2024; Jendrusch and Korbel, 2025), respectively. For dimers, we additionally used a Rosetta interface metric to assess interface plausibility (Stranges and Kuhlman, 2013; Goudy et al., 2023). Starting from the predicted structures by AlphaFold3, we performed 200 iterations of FastRelax (Leman et al., 2020) with the ref2015 energy function (Alford et al., 2017) before interface analysis. Raw Rosetta interface energy is interface-size dependent; therefore, we summarized dimer performance using the fraction of designs satisfying the following size-normalized binding-energy criterion:

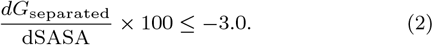

The details of baseline models, metrics, and evaluation settings are provided in the Supplementary Material.

#### 3.1.2. Multi-target binder design

We next evaluated ComplexDesign in a multi-target binder design setting. In this task, a single *de novo* binder is designed to bridge two target proteins within a ternary complex by forming two binder–target interfaces.

We constructed the benchmark from MG-PDB (Liao et al., 2025), a curated dataset of ternary protein complexes bridged by noncovalent molecular glues. To obtain a non-redundant benchmark for this task, we first retained only target pairs in which both target chains were 60–200 residues long. This filter focused the benchmark on medium-sized targets that remain computationally tractable while excluding peptide-like chains. We then reduced redundancy using MMseqs2 (Steinegger and Soding, 2017). Finally, we removed target pairs with high AF3 ipTM for the binary target complex (Abramson et al., 2024), so that the benchmark emphasizes cases not already predicted to form high-confidence binary complexes in the absence of the molecular glue. The resulting benchmark comprises 10 target pairs with target-pair ipTM values ranging from 0.10 to 0.47 (mean 0.273) and spans diverse biological contexts and application settings. The details of benchmark construction and a summary of target pairs are provided in the Supplementary Material.

For each target pair, we performed 1,000 independent binder-design attempts, with binder lengths sampled randomly from the range [10, 110]. Designs that passed the confidence and structural filters were subsequently redesigned with ProteinMPNN (Dauparas et al., 2022), yielding eight binder sequences per design for downstream evaluation. For AlphaFold3 evaluation, we provided MSA and template information for the two target chains and used single-sequence input for the designed binder, following recent binder-design evaluation workflows (Bennett et al., 2023; Latent Labs Team, 2025; Stark et al., 2025). The main metrics reported below are overall AlphaFold3 ipTM and ternary scRMSD.

Here, we do not list the performance of baseline methods for multi-target binder design because, to our knowledge, no existing method directly addresses this setting, which requires one *de novo* binder to engage two separate targets without a predefined target– target arrangement.

### 3.2. ComplexDesign achieves high success rate in designing dimers, trimers, and tetramers

ComplexDesign outperformed the baseline methods in unconditional multichain design, particularly for longer and higher-order assemblies. Across the five dimer settings, ComplexDesign generally achieved higher success rates than APM and Chroma for both original sequences and ProteinMPNN redesigns (Fig. 2a). ProteinMPNN redesign improved APM across several settings, but APM success rates decreased substantially as chain length increased. Chroma rarely met the joint success criteria, achieving only limited success in the shortest length setting.

**Fig. 2:**
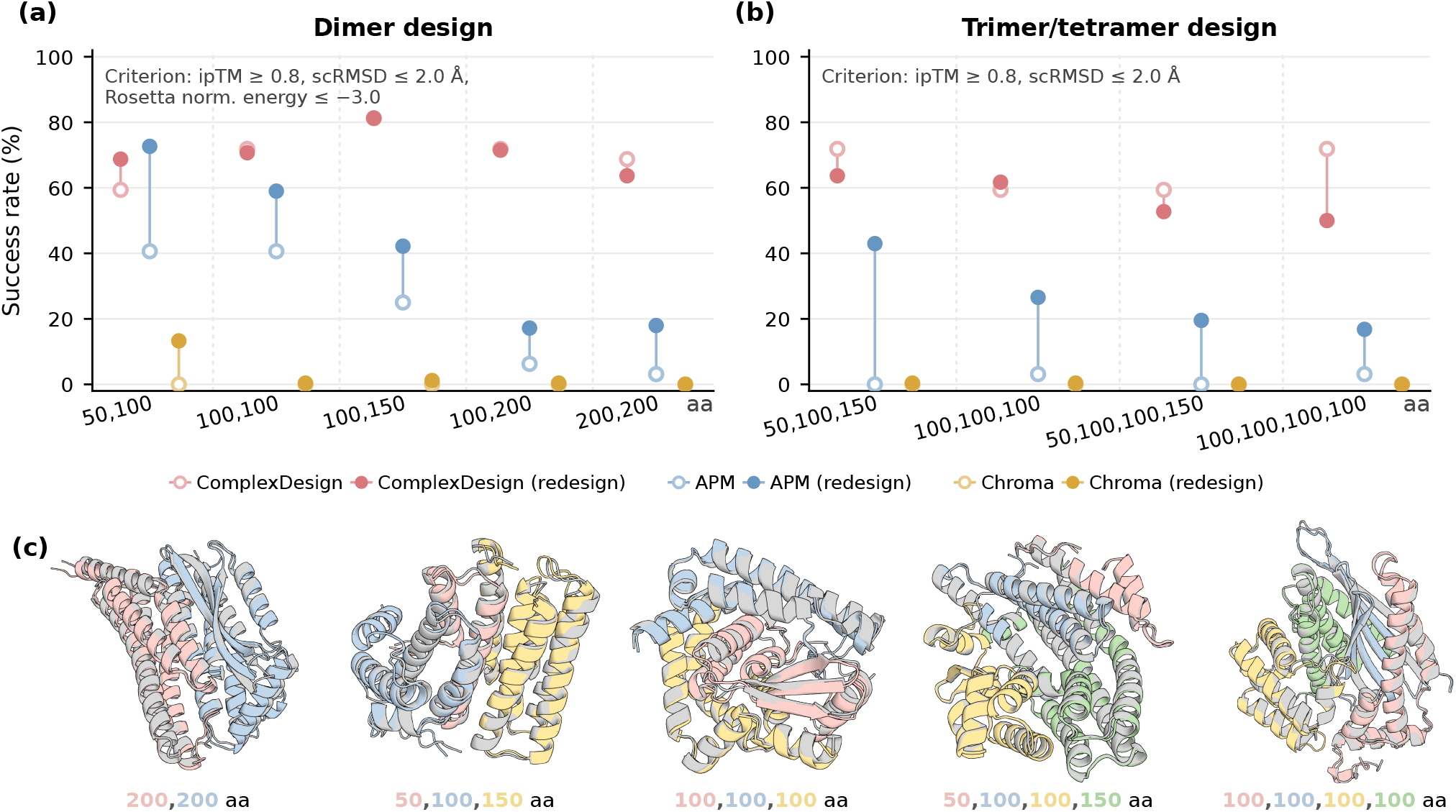
ComplexDesign’s performance in unconditional multichain design. (a,b) Success rates for designs satisfying all criteria indicated in each panel across dimer and trimer/tetramer chain-length settings. Open circles denote original generated sequences, filled circles denote ProteinMPNN redesigns, and vertical lines connect the two values for each method and setting. (c) Representative complexes designed by ComplexDesign. AF3-predicted complexes are shown in color, and the corresponding designed structures are shown in gray after alignment over the full complex. All displayed examples satisfy the corresponding success criteria. **Alt text:** Performance of ComplexDesign in unconditional multichain design. Panels (a) and (b) compare ComplexDesign, APM, and Chroma before and after ProteinMPNN redesign across dimer, trimer, and tetramer design settings. ComplexDesign generally achieves higher success rates than both baselines, with a more pronounced advantage for longer chains and assemblies containing more chains. Panel (c) shows five representative ComplexDesign designed structures aligned with their AlphaFold3 predicted structures.

ComplexDesign showed considerable superiority for higher-order assemblies (Fig. 2b). Across all trimer and tetramer settings, it maintained substantial success rates before and after redesign, whereas APM showed limited success and Chroma remained close to zero. These results indicate that ComplexDesign more reliably supports unconditional design as the number of chains increases.

Fig. 2c shows representative complexes designed by ComplexDesign, which satisfy critical success criteria: AF3 ipTM values of 0.89– 0.94 and C*α* scRMSD values of 0.78–2.00 Å. Single-metric success rates for all three methods are provided in the Supplementary Material.

### 3.3. ComplexDesign yields self-consistent multi-target binders with high confidence

ComplexDesign successfully designed high-confidence ternary complexes for most target pairs in the multi-target binder benchmark. We focus on highly self-consistent designs with ternary scRMSD ≤ 2.0 Å. Figure 3a compares, for each target pair, the AF3 ipTM of the target pair alone with that of the highest-ipTM ternary design satisfying ternary scRMSD ≤ 2.0 Å. Across the benchmark, the target pair alone showed low AF3 confidence.

**Fig. 3:**
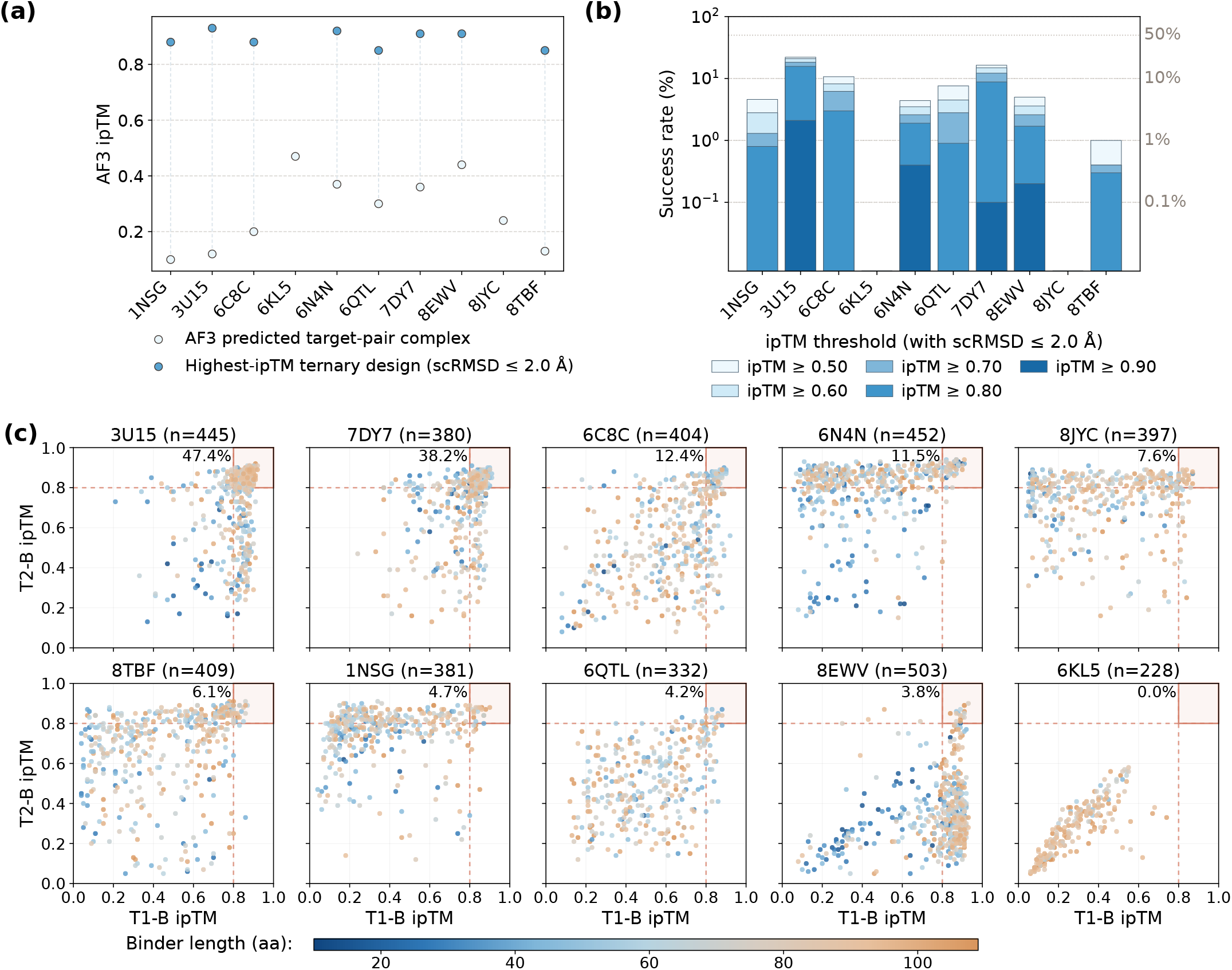
ComplexDesign’s performance in multi-target binder design. (a) AF3 overall ipTM for each target pair before and after binder design. Light blue indicates the target pair alone; dark blue indicates the highest-ipTM ternary design satisfying C*α* scRMSD ≤ 2.0 Å. (b) Per-target success rates across 1,000 independent binder-design attempts, evaluated at ipTM thresholds of 0.50, 0.60, 0.70, 0.80, and 0.90 with ternary scRMSD ≤ 2.0 Å. A design attempt is considered successful if at least one of its eight ProteinMPNN redesigns meets both the scRMSD and ipTM criteria. The y-axis is shown on a logarithmic scale. Darker colors indicate more stringent ipTM thresholds. (c) Interface-confidence distributions of retained designs. Each point represents the highest-ipTM ProteinMPNN redesign from one retained design run. Axes show AF3 interface-specific ipTM values for the binder with target 1 (T1–B) and target 2 (T2–B). Point color indicates binder length. Percentages indicate the fraction of retained designs with both interface-specific ipTM values ≥ 0.8. **Alt text:** Performance of ComplexDesign in multi-target binder design. Panel (a) shows that the target pairs alone have AlphaFold3 overall ipTM values of 0.10–0.47. For the eight target pairs with a ternary design satisfying scRMSD ≤ 2.0 Å, the highest-ipTM designs have ipTM values of 0.85–0.93. Panel (b) shows per-target success rates for designs satisfying scRMSD ≤ 2.0 Å and five different ipTM thresholds. Eight target pairs retain nonzero success rates at ipTM ≥ 0.80, and four remain successful at ipTM ≥ 0.90. Panel (c) shows interface-confidence distributions of retained designs. Successful target pairs such as 3U15 and 7DY7 contain many designs with high confidence at both binder–target interfaces, whereas several other target pairs show asymmetric interface-confidence distributions.

Nevertheless, ComplexDesign produced self-consistent ternary designs for 8 out of the 10 target pairs. Among these successful target pairs, the highest overall AF3 ipTM ranged from 0.85 to 0.93.

Figure 3b summarizes per-target success rates across 1,000 binder-design attempts. A design attempt was considered successful if at least one of its eight ProteinMPNN redesigns met both the scRMSD criterion and the specified ipTM threshold. Notably, eight target pairs showed nonzero success rates at ipTM ≥ 0.8, and four remained successful at the more stringent threshold of ipTM ≥ 0.9. Success varied markedly across targets. The strongest performance was observed for 3U15 and 7DY7, whereas 6KL5 and 8JYC yielded no successful designs under these criteria. For 6KL5, failure was associated with incorrect AF3 prediction of the target structure. For 8JYC, failure was driven by high local RMSD from a long flexible terminal region. Detailed analyses of these two failures are provided in the Supplementary.

Successful designs generally showed balanced confidence and favorable interface features at both binder-target interfaces. We analyzed designs satisfying overall ipTM ≥ 0.8 and scRMSD ≤ 2.0 Å. For 7 of the 8 successful target pairs, the two binder– target interfaces showed similar confidence predicted by AF3: the mean absolute difference between the two interface-specific ipTM values was below 0.05 (Supplementary Table S4). These designs also showed favorable Rosetta interface energies and substantial hydrophobic burial at both binder–target interfaces (Supplementary Tables S5-S7).

### 3.4. Interface-confidence landscapes reveal target-dependent design bottlenecks

Figure 3c shows the retained-design interface-confidence distributions for the 10 target pairs. Retained designs are designs that passed the confidence and structural filters and were carried forward to ProteinMPNN redesign and AF3 evaluation. Each point represents the best of the eight ProteinMPNN redesigns from one retained design run, ranked by overall AF3 ipTM. The axes show the AF3 interface-specific ipTM values for the binder with target 1 (T1–B) and target 2 (T2–B).

The distributions differed markedly across target pairs. 3U15 and 7DY7 contained many retained designs for which both interface-specific ipTM ≥ 0.8, consistent with their high success rates in Figure 3b. In contrast, 1NSG, 6N4N, 8JYC, and 8EWV showed clear imbalance between the two interfaces, with one interface often having higher confidence than the other. These cases suggest that optimization may become trapped in one-sided local optima, but fails to establish a compatible high-confidence interface with the other.

Target properties and binder length also influenced design difficulty. The three target pairs with the highest success rates were all homodimerization cases and generally showed more balanced interface-specific confidence. In contrast, several heterodimerization cases showed more asymmetric interface-confidence distributions. In the heterodimeric cases 1NSG and 6N4N, the lower-confidence interface also mapped to the target surface with higher coil and beta content, suggesting that such surfaces may be harder for the current design procedure to engage reliably. Binder length also mattered: very short binders were rare among retained designs and seldom achieved high confidence at both interfaces.

### 3.5. ComplexDesign shows applicability in various modes of bridging

Representative designs showed that ComplexDesign can design binders to bridge targets through different binder geometries and interface chemistries. We examined the highest-AF3-ipTM cases for 1NSG and 7DY7 (Fig. 4). These target pairs come from distinct biological settings: the FKBP12–FRB(mTOR) target pair derived from the rapamycin-induced ternary complex (Liang et al., 1999) and the PD-L1 target pair derived from a small-molecule-induced PD-L1 dimer (Wang et al., 2022).

**Fig. 4:**
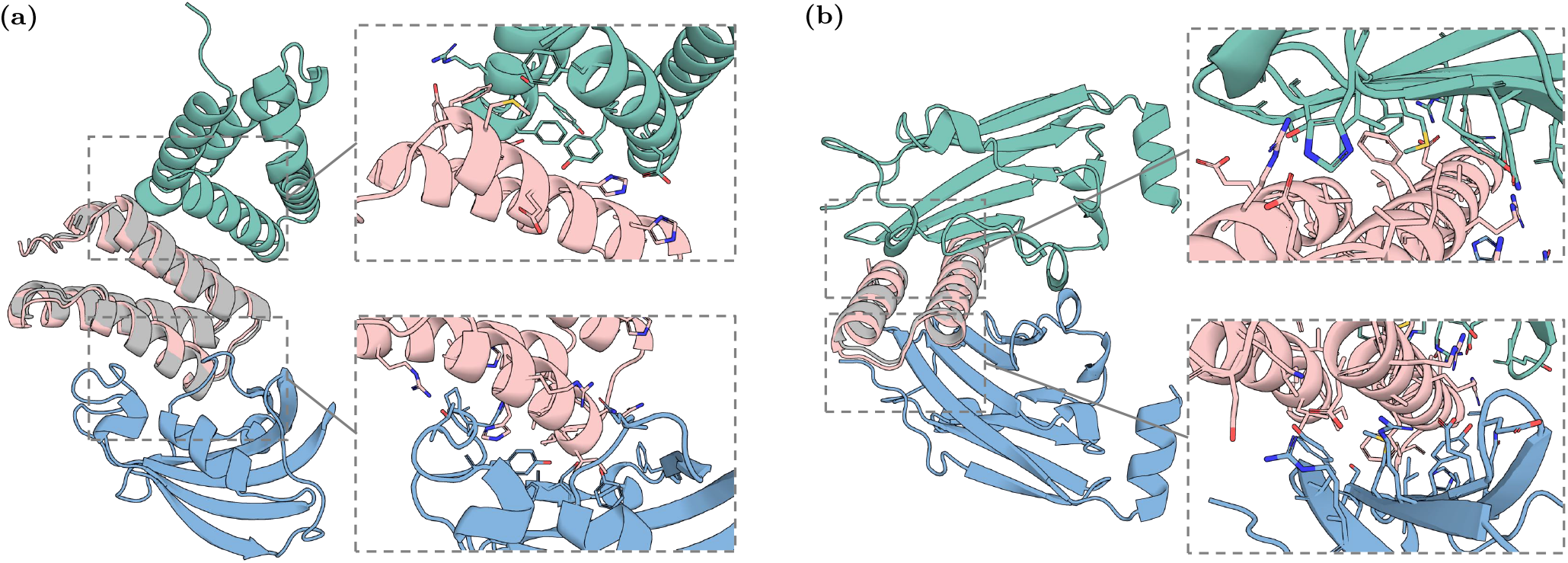
Representative binders designed to bridge two target proteins using ComplexDesign. (a) 1NSG highest AF3-ipTM design (ipTM = 0.88; scRMSD = 0.67 Å). (b) 7DY7 highest AF3-ipTM design (ipTM = 0.91; scRMSD = 1.20 Å). In each panel, the AF3-predicted ternary complex is shown in color, with the binder in pink and the two target chains in teal and blue; the corresponding designed binder structure is shown in gray after full-complex alignment. Dashed boxes mark the two binder–target interfaces, which are enlarged on the right with interface side chains shown as sticks. **Alt text:** Two representative ternary complexes designed by ComplexDesign. Panel (a) shows an 82-residue four-helix binder bridging the FKBP12 and FRB targets in the 1NSG case. Panel (b) shows a compact 43-residue two-helix binder bridging two PD-L1 protomers in the 7DY7 case. In both panels, the designed binder closely aligns with the AlphaFold3-predicted binder structure.

In the case of 1NSG, ComplexDesign produced an 82-residue binder with an AF3-predicted ternary complex ipTM of 0.88 and a scRMSD of 0.67 Å (Fig. 4a). The binder adopts a four-helix architecture that bridges the two targets. The two binder–target interfaces are clearly differentiated. One interface is relatively compact and enriched in hydrophobic packing, whereas the other has a broader contact footprint with hydrophobic packing and a greater contribution from polar contacts.

ComplexDesign also accomplishes successful design in the case of 7DY7: it produced a 43-residue binder with an ipTM of 0.91 and a scRMSD of 1.20 Å (Fig. 4b). In contrast to the 1NSG design, this binder adopts a compact two-helix conformation and packs between the two PD-L1 protomers. It forms dense contacts at both interfaces, dominated by polar and ionic interactions together with local hydrophobic packing.

## 4. Discussion

This study formulates multi-target binder design as a distinct protein complex design problem and introduces ComplexDesign as an initial computational framework for this scenario. In this task, the goal is to design a *de novo* binder that engages two distinct target proteins and forms a plausible ternary complex, in a setting where both the relative arrangement of the targets and the binder interaction sites are unknown. ComplexDesign addresses this challenge by masking pairwise information between the two target chains during optimization. This mechanism avoids predefining the relative geometry of the target chains, providing flexibility for their relative arrangements. This allows the binder– target interfaces and the final ternary arrangement to emerge during design.

Designing multi-target bridging binders requires two coupled capabilities: the binder must adopt a well-defined fold and form productive interfaces with multiple protein partners. The unconditional multichain design results provide a baseline validation of these requirements. Across dimer, trimer, and tetramer designs, ComplexDesign produced complexes with higher predicted confidence and greater structural self-consistency than representative generative baseline methods. The advantage was especially clear for higher-order assemblies, where successful design requires multiple chains to fold and assemble coherently. These results suggest that structure-prediction-guided hallucination is well-suited to multichain design problems that require simultaneous optimization of individual chain folding and inter-chain organization.

Building on this multichain design capability, the multi-target binder benchmark evaluated ComplexDesign for designing binders that bridge two target proteins. The target pairs alone were not predicted by AlphaFold3 to form confident binary complexes, consistent with the intended scenario in which two proteins do not stably associate in the absence of a bridging binder. ComplexDesign nevertheless yielded high-confidence, self-consistent ternary complex designs for most target pairs.

Interface confidence analysis further indicates that the central challenge in multi-target binder design is the coordinated optimization of two binder–target interfaces. When this coordinated optimization fails, the yielded designs often degenerate to one-sided solutions, in which the binder forms an interface with only one target. In fact, this coordination is shaped by both target properties and binder length. First, homodimerization-related target pairs generally show more balanced interface confidence and higher success rates than heterodimeric pairs, suggesting that target similarity or interface symmetry can reduce the difficulty of optimization. Second, targets with surface regions enriched in coil or beta structure, particularly coil-rich surfaces, typically exhibit lower design performance, possibly because these surfaces offer fewer regular interaction geometries than helical surfaces. Third, binder length further modulates designability: very short binders rarely form two high-confidence interfaces, as their limited residue budget makes it more difficult to maintain a stable fold while simultaneously satisfying two binding interfaces.

The present study still has several limitations, which also present opportunities for future research. First, this study focuses on the two-target setting. The same chain-encoding and inter-target masking strategy could be extended to more than two targets, but such higher-order bridging designs remain to be systematically evaluated. Second, the current method requires repeated structure prediction during optimization, which increases computational cost as the number and length of chains grow. Third, our analysis showed that very short binders were underrepresented among high-confidence designs. This suggests a need for more effective strategies for designing short bridging binders. Finally, this study is limited to computational evaluation. The *in silico* metrics used here can help prioritize candidate designs, but experimental characterization will be required to determine whether selected high-confidence sequences can be expressed, fold as predicted, bind their intended targets, and form the designed multichain or ternary complexes.

Together, these results establish ComplexDesign as an effective computational tool for multichain protein design and, more specifically, for designing binders that bridge two target proteins without prescribing their relative arrangement. We anticipate that ComplexDesign will facilitate the development of *de novo* bridging binders, protein-based proximity-inducing systems, and programmable multi-protein assemblies.

## 5. Acknowledgments

The numerical calculations in this study were supported by ICT Computer-X center. We utilized AI tools to assist in grammar correction and translation during the preparation of this manuscript.

## 8. Data availability

The source code of ComplexDesign will be made publicly available upon publication.

## 9. Supplementary data

Supplementary data are available as a separate file associated with this preprint.

