## Supplementary Material for "ComplexDesign: sequence-hallucination design of protein binders bridging multiple proteins"

### 1 Results

#### 1.1 Single-metric evaluation of unconditional multichain designs

To complement the joint success rates shown in Fig. 2, Supplementary Table S1 and S2 report single-metric success rates for unconditional multichain design. Table S1 summarizes dimer results using AF3 overall ipTM, C$\alpha$ scRMSD, and size-normalized Rosetta interface energy as separate criteria, whereas Table S2 summarizes trimer and tetramer results using AF3 overall ipTM and C$\alpha$ scRMSD. Both tables include original generated sequences, and ProteinMPNN redesigns for ComplexDesign, APM, and Chroma.

**Supplementary Table S1:** Evaluation of unconditional dimer design across chain-length settings. Entries report the percentage of designs from ComplexDesign, APM, and Chroma that satisfy each evaluation criterion before and after ProteinMPNN redesign. The criteria are AF3 overall ipTM $\geq 0.8$, C$\alpha$ scRMSD $\leq 2.0$ Å, and normalized Rosetta interface energy $\leq -3.0$. Normalized Rosetta interface energy is defined as $(dG_{\text{separated}}/dSASA) \times 100$.

| | | Chain lengths (aa) | | | | |
|---|---|---|---|---|---|---|
| **Metric** | **Method** | **50,100** | **100,100** | **100,150** | **100,200** | **200,200** |
| **(a) Original sequences** | | | | | | |
| ipTM $\geq 0.8$ (%) | ComplexDesign | **75.00** | **87.50** | **90.62** | **93.75** | **75.00** |
| | APM | 56.25 | 53.12 | 31.25 | 21.88 | 9.38 |
| | Chroma | 18.75 | 3.12 | 0.00 | 0.00 | 0.00 |
| scRMSD $\leq 2.0$ Å (%) | ComplexDesign | **65.62** | **78.12** | **87.50** | **71.88** | **75.00** |
| | APM | **65.62** | 56.25 | 34.38 | 25.00 | 21.88 |
| | Chroma | 18.75 | 0.00 | 3.12 | 0.00 | 0.00 |
| Norm. Rosetta energy $\leq -3.0$ (%) | ComplexDesign | **93.75** | **87.50** | **96.88** | **100.00** | **96.88** |
| | APM | 81.25 | 71.88 | 62.50 | 56.25 | 43.75 |
| | Chroma | 81.25 | 46.88 | 15.62 | 18.75 | 12.50 |
| **(b) ProteinMPNN redesigns** | | | | | | |
| ipTM $\geq 0.8$ (%) | ComplexDesign | **89.06** | **88.28** | **97.66** | **91.80** | **90.23** |
| | APM | 84.77 | 70.70 | 64.06 | 39.84 | 40.23 |
| | Chroma | 21.09 | 3.52 | 4.30 | 1.17 | 0.00 |
| scRMSD $\leq 2.0$ Å (%) | ComplexDesign | 72.27 | **74.61** | **84.77** | **72.66** | **67.58** |
| | APM | **78.52** | 67.97 | 58.98 | 39.84 | 43.36 |
| | Chroma | 22.66 | 1.17 | 2.73 | 1.17 | 0.00 |
| Norm. Rosetta energy $\leq -3.0$ (%) | ComplexDesign | **100.00** | **94.53** | **96.88** | **97.66** | **89.84** |
| | APM | 97.27 | 85.94 | 76.95 | 43.36 | 39.06 |
| | Chroma | 88.28 | 64.06 | 65.23 | 49.22 | 49.22 |

**Supplementary Table S2:** Evaluation of unconditional trimer and tetramer design across chain-length settings. Entries report the percentage of designs from ComplexDesign, APM, and Chroma that satisfy AF3 overall ipTM $\geq 0.8$ or C$\alpha$ scRMSD $\leq 2.0$ Å, before and after ProteinMPNN redesign.

| | | Chain lengths (aa) | | | |
|---|---|---|---|---|---|
| **Metric** | **Method** | **50,100,150** | **100,100,100** | **50,100,100,150** | **100,100,100,100** |
| **(a) Original sequences** | | | | | |
| ipTM $\geq 0.8$ (%) | ComplexDesign | **87.50** | **78.12** | **81.25** | **81.25** |
| | APM | 9.38 | 6.25 | 0.00 | 3.12 |
| | Chroma | 3.12 | 0.00 | 0.00 | 0.00 |
| scRMSD $\leq 2.0$ Å (%) | ComplexDesign | **78.12** | **62.50** | **68.75** | **81.25** |
| | APM | 15.62 | 6.25 | 3.12 | 3.12 |
| | Chroma | 3.12 | 0.00 | 0.00 | 0.00 |
| **(b) ProteinMPNN redesigns** | | | | | |
| ipTM $\geq 0.8$ (%) | ComplexDesign | **92.19** | **89.45** | **80.08** | **82.42** |
| | APM | 62.11 | 37.11 | 35.55 | 28.52 |
| | Chroma | 2.34 | 0.39 | 0.00 | 0.00 |
| scRMSD $\leq 2.0$ Å (%) | ComplexDesign | **64.45** | **62.50** | **53.12** | **51.56** |
| | APM | 46.09 | 35.94 | 23.83 | 21.09 |
| | Chroma | 0.78 | 0.39 | 0.00 | 0.00 |

#### 1.2 Construction of the multi-target binder design benchmark

We constructed the multi-target binder design benchmark from MG-PDB, a curated dataset of 221 ternary complexes in which two protein chains are bridged by a noncovalent molecular glue (Liao et al., 2025).

We first filtered MG-PDB by target-chain length, requiring both target chains to contain 60–200 residues. This criterion excludes peptide-like chains and limits target size, thereby focusing the benchmark on medium-sized proteins that remain computationally tractable for design and evaluation. After this step, 59 candidate complexes remained.

We next reduced redundancy at the target-pair level using MMseqs2 (Steinegger and Söding, 2017). Target-chain sequences were clustered using a minimum sequence identity threshold of 0.4 and a minimum coverage threshold of 0.8 with `-cov-mode 0`. Two target pairs were assigned to the same cluster if any target chain from one pair shared more than 40% sequence identity with any target chain from the other pair. One representative target pair was retained from each cluster, yielding 16 non-redundant target pairs.

We then used AlphaFold3 (Abramson et al., 2024) to predict each target pair as a binary complex and recorded the resulting AF3 ipTM. To focus the benchmark on pairs that are not already predicted to form confident binary complexes in the absence of a bridging partner, we excluded pairs with AF3 ipTM greater than 0.5. The final benchmark comprised 10 target pairs, with AF3 ipTM values ranging from 0.10 to 0.47 (mean 0.273).

The selected targets remain diverse in both biological context and structural characteristics. The benchmark includes infection-related viral systems and several human systems relevant to therapy and immunity. It also spans both homodimerization and heterodimerization settings in the MG-PDB annotation. A target-by-target summary is provided in Supplementary Table S3.

Overall, the resulting benchmark comprises diverse, non-redundant target pairs that are not already predicted to form confident binary complexes without a bridging partner.

**Supplementary Table S3:** Target-by-target summary of the multi-target binder design benchmark.

| PDB | Biological context | MG-PDB annotation | Qualitative structural note | Target-pair ipTM |
|---|---|---|---|---|
| 1NSG | mTOR signaling | Heterodimerization | compact globular pair | 0.10 |
| 3U15 | p53-axis regulation | Homodimerization | symmetric induced dimer | 0.12 |
| 8TBF | RAS-pathway oncology | Heterodimerization | curved interface | 0.13 |
| 6C8C | DNA-damage tolerance | Homodimerization | elongated interface | 0.20 |
| 8JYC | $\gamma\delta$ T-cell immunity | Heterodimerization | flexible terminal region | 0.24 |
| 6QTL | induced proximity / nanobody | Homodimerization | compact domain pair | 0.30 |
| 7DY7 | immune checkpoint | Homodimerization | relatively planar interface | 0.36 |
| 6N4N | antiviral / synthetic control | Heterodimerization | asymmetric globular–helical pair | 0.37 |
| 8EWV | targeted degradation | Heterodimerization | distinct domain sizes | 0.44 |
| 6KL5 | infection-related viral | Homodimerization | loop-rich viral surface | 0.47 |

Biological-context labels provide brief summaries of the biological or application setting of each target pair. The MG-PDB annotation indicates whether the two glue-bridged target proteins are annotated as a homodimerization or heterodimerization case in the original dataset. Qualitative structural notes summarize broad structural characteristics of the extracted target pair relevant to design difficulty. Target-pair ipTM denotes the AlphaFold3-predicted ipTM for the two target chains evaluated without any bridging binder present.

Corresponding target proteins: **1NSG** (FKBP12–FRB/mTOR); **3U15** (MDMX homodimer); **8TBF** (KRAS–CypA); **6C8C** (REV1 CTD–Pol$\kappa$ RIR chimera); **8JYC** (BTN3A1–BTN2A1); **6QTL** (caffeine nanobody homodimer); **7DY7** (PD-L1 homodimer); **6N4N** (DNCR2–NS3A); **8EWV** (VHL–BRD4 BD1); **6KL5** (MERS-CoV N-NTD homodimer).

#### 1.3 Analyses for multi-target binder design

##### 1.3.1 Analysis of successful designs

Successful designs refer to all ProteinMPNN redesigns satisfying overall ipTM $\geq 0.8$ and ternary scRMSD $\leq 2.0$ Å. We analyzed these successful designs from three complementary perspectives: interface-specific confidence, hydrophobicity, and Rosetta interface energy. Table S4 summarizes interface-specific ipTM values. Table S5 reports binder hydrophobic surface exposure together with hydrophobic burial at the two binder–target interfaces. Tables S6 and S7 summarize Rosetta interface energies across successful designs and for the highest-AF3 ipTM successful design of each target pair, respectively.

**Supplementary Table S4:** Per-target summary of interface-specific ipTM values for successful designs.

| Target pair | Successful designs ($n$) | Mean overall ipTM | Mean $T_1$–B ipTM | Mean $T_2$–B ipTM | Mean $\lvert\Delta\rvert$ |
|---|---|---|---|---|---|
| 1NSG | 39 | 0.836 | 0.821 | 0.819 | 0.049 |
| 3U15 | 652 | 0.856 | 0.852 | 0.840 | 0.039 |
| 6C8C | 81 | 0.827 | 0.859 | 0.850 | 0.025 |
| 6N4N | 82 | 0.865 | 0.843 | 0.890 | 0.049 |
| 6QTL | 18 | 0.818 | 0.818 | 0.812 | 0.044 |
| 7DY7 | 332 | 0.834 | 0.831 | 0.836 | 0.039 |
| 8EWV | 59 | 0.859 | 0.892 | 0.843 | 0.049 |
| 8TBF | 9 | 0.822 | 0.730 | 0.872 | 0.142 |

$\lvert\Delta\rvert$ denotes the mean absolute difference between the two interface-specific ipTM values.

As shown in Table S4, across target pairs with successful designs, the two binder–target interfaces usually showed similar confidence. For 7 of the 8 target pairs, the mean absolute difference between the two interface-specific ipTM values was below 0.05. The only exception was 8TBF, for which the two interfaces showed a larger confidence difference.

Table S5 shows that successful designs combined moderate binder RHSA (0.294–0.423) with

**Supplementary Table S5:** Hydrophobic surface and interface burial statistics for successful designs.

| Target pair | Successful designs ($n$) | Binder RHSA | $T_1$–B hyd. bSASA | $T_2$–B hyd. bSASA | $T_1$–B hyd. frac. | $T_2$–B hyd. frac. |
|---|---|---|---|---|---|---|
| 1NSG | 39 | 0.382 | 407.26 | 481.91 | 0.471 | 0.491 |
| 3U15 | 652 | 0.423 | 651.16 | 610.45 | 0.592 | 0.581 |
| 6C8C | 81 | 0.323 | 488.29 | 461.04 | 0.496 | 0.499 |
| 6N4N | 82 | 0.380 | 494.91 | 559.02 | 0.523 | 0.459 |
| 6QTL | 18 | 0.294 | 582.23 | 574.25 | 0.560 | 0.585 |
| 7DY7 | 332 | 0.341 | 463.05 | 484.24 | 0.491 | 0.510 |
| 8EWV | 59 | 0.339 | 608.25 | 534.59 | 0.614 | 0.502 |
| 8TBF | 9 | 0.359 | 408.18 | 384.11 | 0.545 | 0.434 |

RHSA denotes binder relative hydrophobic surface area; hyd. bSASA denotes hydrophobic buried solvent-accessible surface area; hyd. frac. denotes hydrophobic buried fraction. Hydrophobic buried SASA values are reported in $Å^2$; RHSA and hydrophobic buried fraction are unitless.

**Supplementary Table S6:** Rosetta interface energy summary across successful designs.

| Target pair | Successful designs ($n$) | Mean $\Delta G_{T_1\text{-}B}$ | Mean $\Delta G_{T_2\text{-}B}$ | Mean less favorable interface $\Delta G$ |
|---|---|---|---|---|
| 1NSG | 39 | -45.45 | -53.79 | -42.94 |
| 3U15 | 652 | -62.87 | -59.05 | -55.20 |
| 6C8C | 81 | -56.64 | -52.46 | -49.30 |
| 6N4N | 82 | -49.58 | -63.33 | -48.91 |
| 6QTL | 18 | -56.18 | -54.22 | -49.62 |
| 7DY7 | 332 | -50.94 | -51.52 | -47.17 |
| 8EWV | 59 | -67.19 | -49.09 | -48.98 |
| 8TBF | 9 | -35.95 | -48.50 | -35.87 |

$\Delta G_{T_1\text{-}B}$ and $\Delta G_{T_2\text{-}B}$ denote the pairwise binder–target interface energies after all-atom refinement. Mean less favorable interface $\Delta G$ is the mean, across successful designs, of the less favorable (less negative) of the two per-design interface energies. All energies are reported in REU.

**Supplementary Table S7:** Summary of the highest-ipTM successful design for each target pair.

| Target pair | Binder length | ipTM | scRMSD (Å) | $\Delta G_{T_1\text{-}B}$ | $\Delta G_{T_2\text{-}B}$ |
|---|---|---|---|---|---|
| 1NSG | 82 | 0.88 | 0.67 | -40.83 | -61.53 |
| 3U15 | 74 | 0.93 | 0.47 | -89.03 | -66.14 |
| 6C8C | 47 | 0.88 | 1.15 | -67.19 | -37.57 |
| 6N4N | 95 | 0.92 | 1.78 | -62.00 | -62.73 |
| 6QTL | 90 | 0.85 | 1.23 | -64.89 | -51.87 |
| 7DY7 | 43 | 0.91 | 1.20 | -40.98 | -47.09 |
| 8EWV | 77 | 0.91 | 1.44 | -58.89 | -45.65 |
| 8TBF | 47 | 0.85 | 1.86 | -29.42 | -46.99 |

For each target pair, the reported design is the successful design with the highest AlphaFold3 ipTM. Reported values include binder length, overall ipTM, ternary scRMSD, and the two pairwise binder–target interface energies. All Rosetta energies are reported in REU.

substantial hydrophobic burial at both binder–target interfaces. Across target pairs, the mean hydrophobic buried SASA was appreciable on both sides. These summaries are consistent with hydrophobic packing contributing to both interfaces in successful designs, without requiring unusually high overall binder hydrophobicity.

Rosetta analysis further supported the energetic plausibility of these designs. Table S6 reports the two pairwise binder–target interface energies separately. Even when each design was summarized by its less favorable interface, the mean $\Delta G$ remained favorable across all target pairs, ranging from −35.87 to −55.20 REU.

Table S7 summarizes, for each target pair, the successful design with the highest AF3 ipTM. Across all eight target pairs with successful designs, these representatives showed high AF3 confidence (ipTM 0.85–0.93), low ternary scRMSD (0.47–1.86 Å), and favorable Rosetta interface energies on both binder–target interfaces.

##### 1.3.2 Failure analysis of 6KL5 and 8JYC

Two target pairs in the benchmark did not yield successful designs under the main criterion of overall ipTM $\geq 0.8$ and ternary scRMSD $\leq 2.0$ Å. To illustrate their failure reasons, Fig. S1 shows one representative design from each target pair. These two cases exhibit different target-specific failure modes.

For 6KL5, the main issue arises during AlphaFold3 reevaluation (Fig. S1a,b). The AF3-predicted structure does not preserve the target structures seen in the designed complex. Instead, it adopts a markedly different incorrect arrangement in which the two target chains interpenetrate. This target-side prediction failure strongly degrades the full-complex metrics, yielding an all-chain C$\alpha$ scRMSD of 25.05 Å and an overall ipTM of 0.58. Per-chain scRMSDs are also very large for both targets (25.32 and 28.99 Å), and remain substantial for the binder (16.45 Å). These values indicate that failure is driven primarily by incorrect target prediction, including both target structure and target–target arrangement. As a result, the designed binder-mediated ternary geometry is not recovered, and AF3 assigns low overall confidence to the complex.

A different failure mode is observed for 8JYC (Fig. S1c). The representative design has a high overall AF3 ipTM (0.86) but fails under the scRMSD criterion. The elevated scRMSD is mainly caused by a flexible terminal segment in target 1. In the designed complex, this segment follows the crystal-structure template and extends outward. In the AF3-predicted model, the same segment folds back onto the surface of target 1, producing a large local deviation beginning at residue A182. As a result, the all-chain C$\alpha$ RMSD increases to 5.06 Å, even though the binder and target 2 remain well aligned, with C$\alpha$ RMSDs of 1.82 and 1.75 Å, respectively. When the high-deviation segment (A182–A200) is excluded, the ternary C$\alpha$ RMSD decreases to 1.31 Å, below the scRMSD success threshold. This result indicates that the failure is dominated by a localized mismatch in a flexible terminal region rather than by a global failure of ternary-complex formation.

Together, these cases suggest that the remaining failures in this benchmark are target-specific. They also show that, under AF3-based evaluation with a strict full-complex scRMSD criterion, target-side prediction errors or flexible-region deviations can cause otherwise plausible ternary designs to fall outside the success threshold.

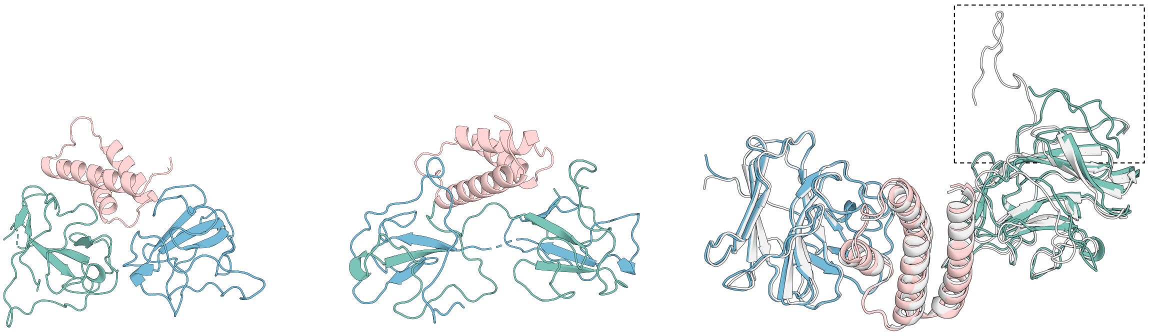

**(a) 6KL5: designed complex** **(b) 6KL5: AF3-predicted complex** **(c) 8JYC: alignment of designed and AF3-predicted complexes**

**Supplementary Fig. S1:** Representative failure modes for 6KL5 and 8JYC in the multi-target binder design benchmark. In all panels, target 1 is shown in green, target 2 in blue, and the binder in pink; in panel (c), the designed complex is shown in gray and the AF3-predicted complex in color. (a,b) For 6KL5, the AF3-predicted complex shows severe target-chain interpenetration and large deviation from the designed ternary geometry. (c) For 8JYC, the boxed region highlights a flexible terminal segment in target 1 that accounts for most of the structural deviation. Additional quantitative analysis is provided in Supplementary Section 1.3.2.

### 2 Methods

#### 2.1 Loss terms used in the optimization objective

The optimization objective used in this work follows the implementations in ColabDesign and BindCraft (Ovchinnikov et al., 2025; Pacesa et al., 2025). This section summarizes the function, weight, and application scope of each loss term in the two design tasks.

- **pLDDT (weight 0.1).** This term encourages high-confidence local structure. In unconditional multichain design, it is applied to all residues. In multi-target binder design, it is applied to binder residues only.

- **pAE (weight 0.4).** This term penalizes large predicted aligned errors and thus favors more reliable relative residue placement. In unconditional multichain design, it is computed over same-chain residue pairs. In multi-target binder design, it is computed over residue pairs involving the binder, including both binder–binder and binder–target pairs.

- **Contact (weight 1.0).** This term encourages predicted residue–residue contacts and thus promotes compact packing. In unconditional multichain design, it is computed over same-chain residue pairs, excluding pairs with sequence separation smaller than 9. In multi-target binder design, it is computed over binder–binder residue pairs, also excluding pairs with sequence separation smaller than 9.

- **i_pAE (weight 0.1).** This interface-related term penalizes large predicted aligned errors between chains. In unconditional multichain design, it is computed over inter-chain residue pairs. In multi-target binder design, it is computed over binder–target residue pairs.

- **i_contact (weight 1.0).** This interface-specific term encourages predicted residue–residue contacts between chains. In unconditional multichain design, it is computed over inter-chain residue pairs. In multi-target binder design, it is computed over binder–target residue pairs.

- **i_pTM (weight 0.05).** This term encourages higher predicted interface TM scores and thus favors greater confidence in the overall inter-chain arrangement. In both tasks, it is computed using all different-chain residue pairs in the predicted complex.
- **Radius of gyration (weight 0.3; multi-target binder design only).** This term is used only in multi-target binder design. It penalizes overly extended binder conformations and encourages compact folded states.

#### 2.2 Baseline methods and design settings

For the unconditional multichain design benchmark, we compared ComplexDesign with two representative baseline methods, APM (Chen et al., 2025) and Chroma (Ingraham et al., 2023). Both baselines were evaluated under the same nine chain-length combinations as ComplexDesign: $[50, 100]$, $[100, 100]$, $[100, 150]$, $[100, 200]$, $[200, 200]$, $[50, 100, 150]$, $[100, 100, 100]$, $[50, 100, 100, 150]$, and $[100, 100, 100, 100]$.

For APM, we used the official unconditional multimer inference script with the released refinement checkpoint. We changed only the sampling settings needed to match our benchmark. Specifically, we set `samples_per_length` to 32 and `num_batch` to 1, and restricted generation to the nine chain-length combinations listed above. All other settings followed the default configuration of the official implementation.

For Chroma, we used the official Python API together with a custom batch-sampling script. The script repeatedly called `chroma.sample(chain_lengths=lengths)` for each of the same nine chain-length combinations. For each setting, we generated 32 complexes using the default sampling behavior of the official API.

All complexes generated by ComplexDesign, APM, and Chroma were evaluated using the same downstream pipeline. We evaluated both the original generated sequences and the corresponding ProteinMPNN redesigns (Dauparas et al., 2022). We then applied the same AlphaFold3-based evaluation (Abramson et al., 2024), scRMSD calculation, and, for dimers, size-normalized Rosetta interface analysis to all methods.

#### 2.3 ProteinMPNN redesign settings

For ProteinMPNN redesign (Dauparas et al., 2022), we generated eight redesigned sequences for each structure using the original ProteinMPNN weights with model `v_48_020`, a sampling temperature of 0.1, and no backbone noise. In the unconditional multichain design task, all chains were redesigned. In the multi-target binder design task, binder residues with at least one atom within 3.5 Å of either target were kept fixed, whereas all other binder residues were allowed to vary during redesign.

#### 2.4 Additional evaluation details

##### 2.4.1 Unconditional multichain design

For unconditional multichain design, all generated complexes were evaluated with AlphaFold3 in single-sequence mode, with no MSA or template information provided for any chain (Abramson et al., 2024). The reported ipTM is the overall ipTM of the AlphaFold3-predicted complex.

The scRMSD was computed as the $C_\alpha$ RMSD between the AlphaFold3-predicted complex and the corresponding generated structure after global alignment over all chains.

For the dimer benchmark, we additionally evaluated pairwise Rosetta interface energetics on the AlphaFold3-predicted complexes. Before scoring, each structure was relaxed for 200 iterations using FastRelax (Leman et al., 2020) with the ref2015 energy function (Alford et al., 2017). Interface properties were then computed with InterfaceAnalyzerMover (Stranges and Kuhlman, 2013) with separated-state repacking enabled. We summarized dimer interface energetics using the size-normalized score $(dG_{\text{separated}}/dSASA) \times 100$, where $dG_{\text{separated}}$ is the Rosetta separated binding energy and $dSASA$ is the buried solvent-accessible surface area of the interface. This normalization reduces the dependence of raw $dG_{\text{separated}}$ on interface size. Rosetta documentation describes values below $-1.5$ for this metric as generally favorable (RosettaCommons, 2025). In our dimer benchmark, however, nearly all dimer designs produced by three methods satisfied this threshold. It therefore provided little discrimination between methods. We therefore used the more stringent cutoff of $-3.0$ for comparative evaluation.

##### 2.4.2 Multi-target binder design

For multi-target binder design, the two target chains were provided to AlphaFold3 with MSAs and structural templates from the standard data pipeline. No MSA or template information was provided for the designed binder, consistent with recent binder-design evaluation settings (Bennett et al., 2023; Li et al., 2025). Unless otherwise specified, ipTM refers to the overall AlphaFold3 ipTM of the full ternary complex. scRMSD was computed as in the unconditional multichain design task, using all $C_\alpha$ atoms from all three chains after global alignment.

For supplementary analyses of successful designs, we evaluated the two binder–target interfaces separately. Here, $T_1$ and $T_2$ denote the two target chains, and B denotes the designed binder. Interface-specific ipTM values for the $T_1$–B and $T_2$–B interfaces were obtained from the AlphaFold3 chain-pair ipTM outputs for the predicted ternary complex.

We also computed pairwise Rosetta interface energies separately for the $T_1$–B and $T_2$–B interfaces using the same relaxation and InterfaceAnalyzerMover protocol as in the unconditional multichain design task. In contrast to the size-normalized Rosetta metric used for the unconditional dimer benchmark, we report raw Rosetta interface energy here as a measure of interface quality, following prior binder-design studies (Cao et al., 2022; Bennett et al., 2023).

We further quantified hydrophobicity using two SASA-based metrics. At the binder level, binder relative hydrophobic surface area (RHSA) was defined as the fraction of total binder SASA contributed by hydrophobic residues. At the interface level, the hydrophobic buried fraction was computed separately for the $T_1$–B and $T_2$–B interfaces, defined as the fraction of buried interface SASA contributed by hydrophobic residues. Hydrophobic residues were defined as Ala, Val, Ile, Leu, Met, Phe, Trp, Tyr, and Pro.
